# Reduced Nicotinamide Mononucleotide (NMNH) Potently Enhances NAD^+^, Suppresses Glycolysis, TCA Cycle and Cell Growth

**DOI:** 10.1101/2020.11.03.366427

**Authors:** Yan Liu, Chengting Luo, Ting Li, Wenhao Zhang, Zhaoyun Zong, Xiaohui Liu, Haiteng Deng

## Abstract

In the present study, we developed a chemical method to produce dihydro nicotinamide mononucleotide (NMNH), which is the reduced-form of nicotinamide mononucleotide (NMN). We demonstrated that NMNH was a better NAD^+^ enhancer compared to NMN both *in vitro* and *in vivo* mediated by NMNAT. Additionally, NMNH increased the reduced NAD (NADH) levels in cells and in mouse liver. Metabolomic analysis revealed that NMNH inhibited glycolysis and TCA cycle. *In vitro* experiments demonstrated that NMNH induced cell cycle arrest and suppressed cell growth. Nevertheless, NMNH treatment did not cause observable difference in mice. Taken together, our work demonstrates that NMNH is a potent NAD^+^ enhancer, and suppresses glycolysis, TCA cycle and cell growth.

## INTRODUCTION

Nicotinamide adenine dinucleotide (NAD^+^) is essential to living organisms because it participates in hundreds of biological reactions and regulates key biological processes, such as metabolism and DNA repair.^1, 2^ Previous studies revealed that upregulating NAD^+^ biosynthesis by genetic manipulation leads to increased stress resistance and elongated lifespan in yeast and *Drosophila.*^3–5^ Increasing NAD^+^ level is proved to delay progeroid and other degenerative disease in mice.^6, 7^ There is ample evidence that nicotinamide riboside (NR) and nicotinamide mononucleotide (NMN) are potent NAD^+^ enhancers, as they increase cellular NAD^+^ levels and confer multiple health benefits.^8^ Supplementation with NR or NMN protects mice against age-associated health deterioration. ^9, 10^

These studies mainly focused on NAD precursors in the oxidized form since most NAD^+^ consuming enzymes uses NAD^+^ as the substrate. Less is known about the roles of NAD precursors in the reduced form. Two recent studies revealed that NR in its reduced form, denoted as NRH, was a better NAD^+^ booster than NR or NMN in cells and tissues.^11, 12^ Yang *et al.* also showed that NRH increased resistance to cell death caused by genotoxins and the conversion of NRH to NMNH was independent of Nrk1 or Nrk2. Giroud-Gerbetant et al. found that NRH was converted to NMNH by adenosine kinase to synthesize NAD^+^, and demonstrated that NRH was orally available and prevented cisplatin-induced acute kidney injury in mice. However, it has not been reported, to the best of our knowledge, whether the reduced form of NMN can increase cellular NAD^+^ levels and regulate biological processes.

In the present work, we developed a chemical reduction method for synthesizing the reduced NMN, denoted as NMNH and investigated the biological effects of NMNH on cellular processes. We found that NMNH was a better NAD^+^ enhancer than NMN both *in vitro* and *in vivo*. Moreover, NMNH increased cellular NADH levels, suppressed glycolysis and TCA cycle, as well as cell growth.

## RESULTS

### NMNH synthesis

We investigated enzymatic and chemical synthesis methods for preparing NMNH and found that NMNH generated from chemical synthesis had a high yield and high purity. The procedure for NMNH synthesis is displayed in Figure 1A. NMNH was produced by reduction of NMN using thiourea dioxide (TDO). We chose TDO as the reducing reagent based on the facts that TDO is stable in common laboratory storage conditions, and it is amenable to solution preparation. TDO possesses high reduction potential in aqueous solutions and is also suitable for scalable applications. Briefly, 340 mg NMN and 125 mg TDO were dissolved in 1 mL of 10% ammonia solution. The reaction mix was incubated at 40°C for 1 hour and the product was then purified by HPLC using an amide column followed by vacuum drying. Purified NMNH was in orange color and had a UV absorption centered at 340 nm (Figure 1B). The reaction product was characterized by high resolution mass spectrometry. The mass to charge ratio (m/z) of the final product was determined to be 335.0648 in negative ion mode, which matches the theoretical molecular weight of NMNH (Figure S1A) with an elemental composition of C11H17N2O8P1. These results suggest that the reduction product is NMNH.

**Figure 1.**
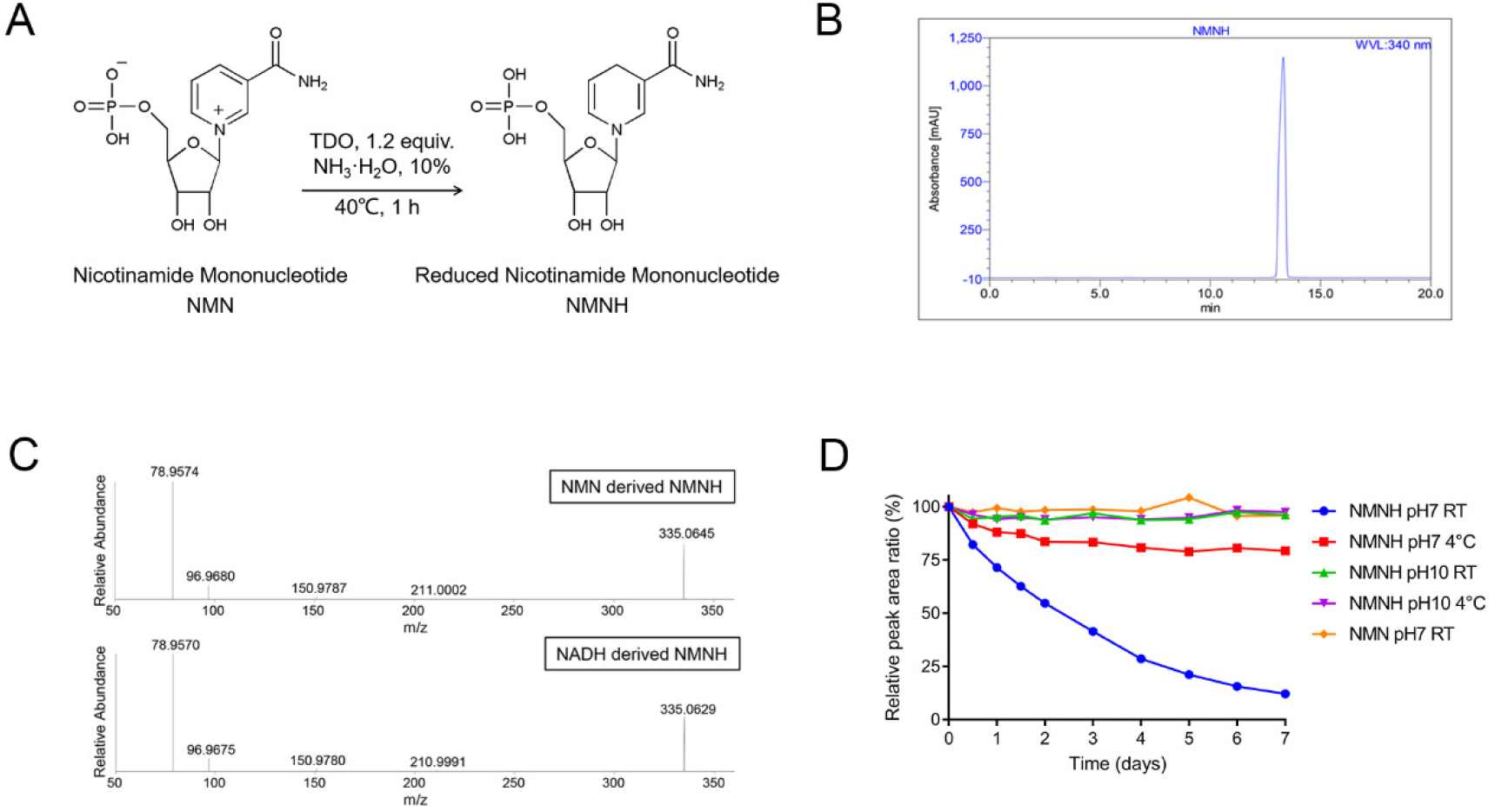
NMNH synthesis. (A) NMNH synthesis procedure. NMNH was generated by reducing NMN with TDO under alkaline environment at 40 °C for 1 h. (B) Characteristic UV absorption peak of NMNH. NMNH has a UV absorption centered at 340 nm. (C) MS/MS spectra comparison of enzymatically- and chemically-generated NMNH. NMNH generated by NMN reduction has the identical fragmentation pattern as NMNH generated by NADH decomposition. (D) NMNH is more stable under alkali conditions and at low temperature. Equal amount of NMNH and NMN were stored under different conditions and their 340 nm and 260 nm absorptions were determined.

To further verify the identity of the reaction product, we generated NMNH from NADH using NADH pyrophosphatase (NudC), which decomposes NADH into NMNH and AMP ^13^. The NADH decomposition reaction is shown in Figure S1B. The NudC-generated product was analyzed using high resolution mass spectrometry and the major peak was observed with mass to charge ratio (m/z) 335.0633 in negative ion mode (Figure S1C), precisely matching the mass of the reduction-generated NMNH. We compared the MS/MS spectra of enzymatically- and chemically-generated NMNH and found that they shared an identical fragmentation pattern (Figure 1C). Therefore, we confirmed that reduction of NMN by TDO produces NMNH. These results show that this reduction method is robust and effective. We examined the stability of NMNH and NMN and found that NMNH was stable under alkaline pH and low temperature conditions (Figure 1D). Unlike NMN, NMNH was unstable at neutral pH in solution (Figure 1D). The half-life of NMNH at pH 7.0 and at room temperature was about 2.4 days. We also found that NMNH was less stable than NMN in cell medium (Figure S1D).

### NMNH increased cellular NAD^+^ levels both *in vitro* and *in vivo*

We next examined the effects of NMNH on enhancing cellular NAD^+^ levels as compared to NMN. NMNH (100 μM) treatment increased the level of cellular NAD^+^ by 5 folds in HepG2 cells whereas 100 μM NMN only slightly increased the level of NAD^+^, as determined by mass spectrometry (Figure 2A). The result indicates that, compared with NMN, NMNH is a more efficient NAD^+^ enhancer. In addition to HepG2 cells, we also confirmed that NMNH enhanced the NAD^+^ in ES-2 cells and 3T3-L1 cells, which are human ovary derived cells and mouse embryo fibroblast derived cells, respectively (Figure S2A and S2B). We found that NMNH increased cellular NAD^+^ levels in time- and concentration-dependent manner in 786-O cells (Figure S2C and S2D). We also found that NMNH increased cellular NAM concentration (Figure 2B).

**Figure 2.**
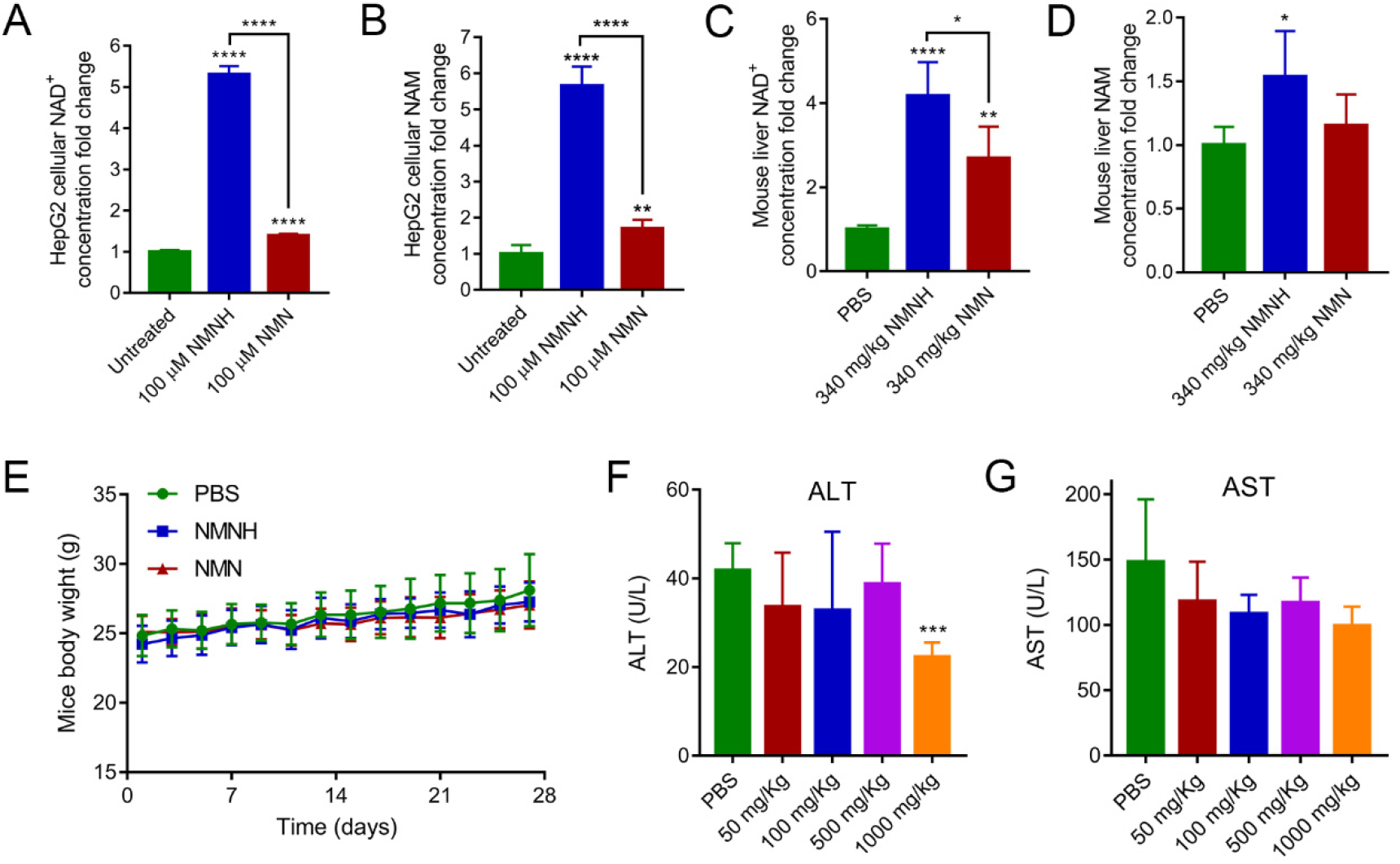
NMNH is potent NAD^+^ booster both *in vitro* and *in vivo*. (A) NMNH had better NAD^+^ enhancing effect than NMN in HepG2 cell. Cellular NAD^+^ concentration was determined after 100 μM NMN or NMNH treatment for 12 h using TSQ Quantiva mass spectrometer. Data are shown as mean ± SD (n = 4). ****p < 0.0001. Significance is obtained compared to the untreated group unless otherwise specified. (B) NMNH increased cellular NAM concentration in HepG2 cell. Cellular NAM concentration was determined after 100 μM NMN or NMNH treatment for 12 h using Q Exactive mass spectrometer. Data are shown as mean ± SD (n = 4). **p < 0.01, ****p < 0.0001. Significance is obtained compared to the untreated group unless otherwise specified. (C) NMNH had better *in vivo* NAD^+^ enhancing effect than NMN. C57BL/6J male mice were treated with 340 mg/kg NMN or NMNH for 6 h by intraperitoneal injection. Liver NAD^+^ concentration was determined using TSQ Quantiva mass spectrometer. Data are shown as mean ± SD (n = 5). *p < 0.05, **p < 0.01, ****p < 0.0001. Significance is obtained compared to the untreated group unless otherwise specified. (D) NMNH increased liver NAM concentration. C57BL/6J male mice were treated with 340 mg/kg NMN or NMNH for 6 h by intraperitoneal injection. Liver NAM concentration was determined using TSQ Quantiva mass spectrometer. Data are shown as mean ± SD (n = 5). *p < 0.05. Significance is obtained compared to the untreated group unless otherwise specified. (E) Mice body weight curves under NMNH or NMN treatment. C57BL/6J male mice were treated with 13.6 mg/kg NMN or NMNH every day for 4 weeks by intraperitoneal injection. Data are shown as mean ± SD (n = 5). (F) Mice serum ALT levels after different dosages of NMNH treatment. C57BL/6J male mice were treated with 50, 100, 500, 1000 mg/kg NMNH or PBS every other day for 1 week by intraperitoneal injection. Data are shown as mean ± SD (n = 5 for PBS, 50 and 100 mg/kg groups; n = 4 for 500 and 1000 mg/kg groups). (G) Mice serum AST levels after different dosages of NMNH treatment. C57BL/6J male mice were treated with 50, 100, 500, 1000 mg/kg NMNH or PBS every other day for 1 week by intraperitoneal injection. Data are shown as mean ± SD (n = 5 for PBS, 50 and 100 mg/kg groups; n = 4 for 500 and 1000 mg/kg groups).

We further examined the effects of NMNH on NAD^+^ levels using a mouse model. We treated C57BL/6J male mice with 340 mg/kg NMN or NMNH via intraperitoneal injection and measured liver NAD^+^ concentrations by mass spectrometry. We found that the liver NAD^+^ level was 4-fold higher in NMNH-treated mice than that in PBS-treated mice (Figure 2C). Additionally, the liver NAD^+^ level in NMNH-treated mice was 1.5-fold higher than that in NMN-treated mice. These results demonstrate that NMNH is a more potent *in vivo* NAD^+^ enhancer than NMN. NMNH also significantly increased levels of NAM in mouse liver (Figure 2D). We also examined the NAD^+^ enhancing effect of NMNH by oral delivery into mice and found that liver NAD^+^ level was increased by nearly 4 folds (Figure S2E).

To verify whether NMNH shows any repression effect in animal, we treated C57BL/6J male mice with 13.6 mg/kg NMN or NMNH via intraperitoneal injection every day for 4 weeks and found no difference in body weight curves among PBS-treated, NMNH- and NMN-treated mice, as shown in Figure 2E.

We also examined whether the high dosage of NMNH induced liver toxicity. We treated C57BL/6J male mice with 50, 100, 500 or 1000 mg/kg NMNH via intraperitoneal injection every other day for a week and found that the serum levels of alanine aminotransferase (ALT) and aspartate aminotransferase (AST) were not elevated (Figure 2F and 2G), which suggested that NMNH was not liver toxic at high dosage. We also found that NMNH treatment increased mouse liver NAD^+^ contents in a dosage-dependent manner, and 1000 mg/kg NMNH treatment increased mouse liver NAD^+^ level by over 10 folds, as shown in Figure S2F.

### NMNH inhibited Glycolysis and TCA Cycle

It has been proposed that NRH is converted to NMNH, which is further converted to NADH by nicotinamide mononucleotide adenylyltransferase (NMNAT).^12, 14^ To examine whether NMNH increases the cellular NADH concentration, we measured NADH using mass spectrometry. We found that NADH levels were 2.5-fold higher in NMNH-treated HepG2 cells than in untreated and NMN-treated cells (Figure 3A). Increased NADH level was also detected in NMNH-treated 786-O cells (Figure S3A). We also uncovered that 340 mg/kg NMNH treatment increased NADH level by 3 folds in mouse liver while 340 mg/kg NMN treatment increased NADH level by 1.7 folds (Figure 3B). Furthermore, we found that NMNH treatment increased mouse liver NADH contents in a dosage-dependent manner, and 1000 mg/kg NMNH treatment increased mouse liver NADH level by nearly 10 folds (Figure S3B).

**Figure 3.**
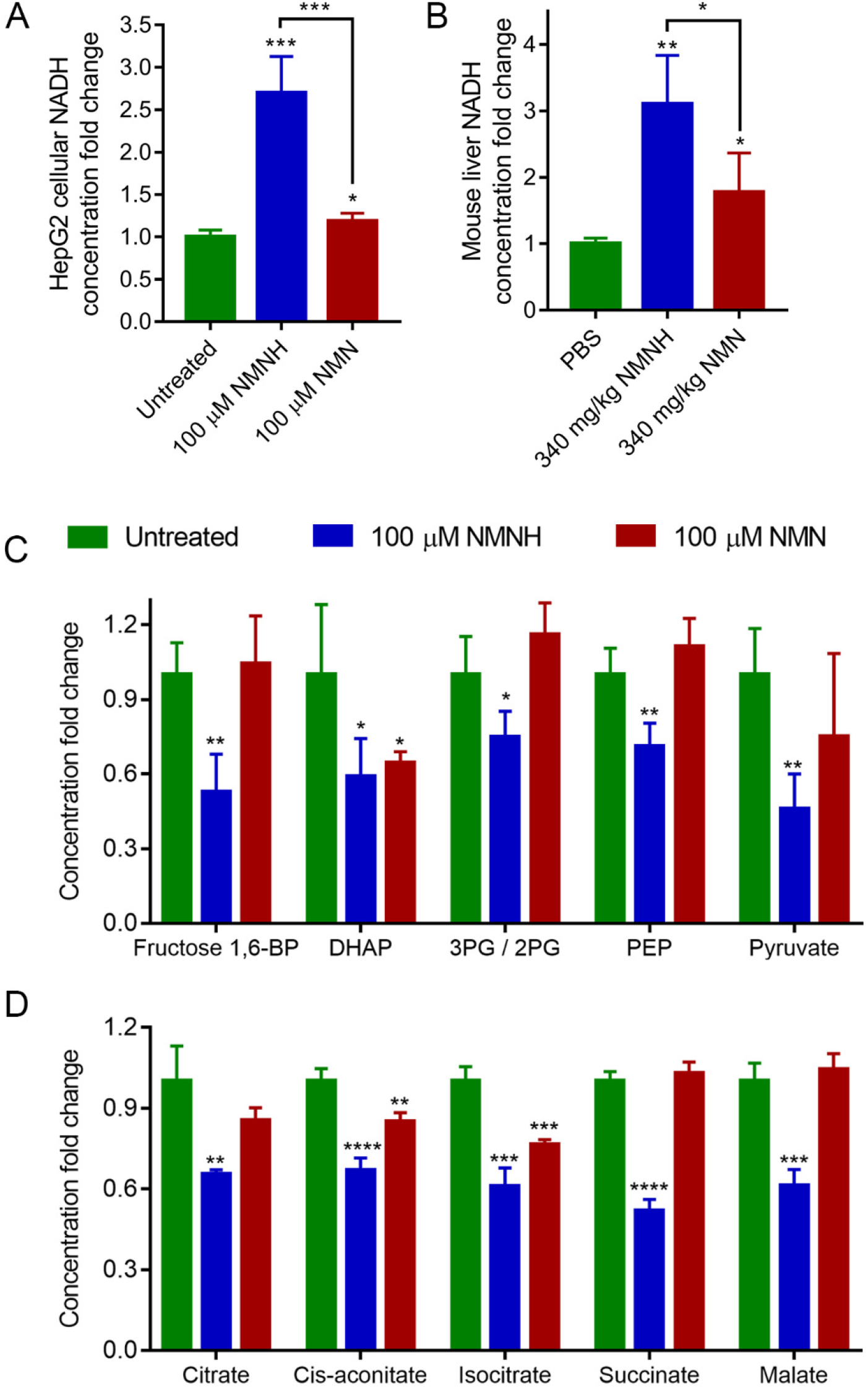
NMNH repressed glycolysis and TCA cycle. (A) NMNH increased cellular NADH concentration in HepG2 cell. Cellular NADH concentration was determined after 100 μM NMN or NMNH treatment for 12 h using TSQ Quantiva mass spectrometer. Data are shown as mean ± SD (n = 4). *p < 0.05, ***p < 0.001. Significance is obtained compared to the untreated group unless otherwise specified. (B) NMNH increased liver NADH concentration. C57BL/6J male mice were treated with 340 mg/kg NMN or NMNH for 6 h by intraperitoneal injection. Liver NADH concentration was determined using TSQ Quantiva mass spectrometer. Data are shown as mean ± SD (n = 4). *p < 0.05, **p < 0.01. Significance is obtained compared to the untreated group unless otherwise specified. (C) NMNH decreased glycolysis intermediate concentrations in HepG2 cell. Cellular metabolite concentrations were determined after 100 μM NMN or NMNH treatment for 12 h using TSQ Quantiva mass spectrometer. Data are shown as mean ± SD (n = 4). *p < 0.05, **p < 0.01. Fructose-1,6-BP, fructose-1,6-bisphosphate. DHAP, dihydroxyacetone phosphate. 3PG / 2PG, 3-phosphoglycerate / 2-phosphoglycerate. PEP, phosphoenolpyruvate. Significance is obtained compared to the untreated group unless otherwise specified. (D) NMNH decreased TCA cycle intermediate concentrations in HepG2 cell. Cellular metabolite concentrations were determined after 100 μM NMN or NMNH treatment for 12 h using TSQ Quantiva mass spectrometer. Data are shown as mean ± SD (n = 4). **p < 0.01, ***p < 0.001, ****p < 0.0001. Significance is obtained compared to the untreated group unless otherwise specified.

Previous studies indicated that accumulation of NADH slowed down glycolysis and TCA cycle.^15, 16^ To examine cellular metabolic responses to NMNH treatment, we carried out metabolomic profiling on NMNH- and NMN-treated cells. We found that NMNH treatment decreased levels of glycolysis intermediates, including fructose-1,6-diphosphate, DHAP, 3PG / 2PG, PEP and pyruvate (Figure 3C). NMNH treatment also decreased the levels of TCA cycle intermediates, including citrate, cis-aconitate, isocitrate, succinate, and malate (Figure 3D). These results indicated that NMNH-generated NADH accumulation inhibits cellular glycolysis and the TCA cycle.

We further carried out isotope tracing experiment using ^13^C_6_-glucose to trace the regulation of glycolysis and TCA cycle by NMNH. As shown in Figure S3C, intermediates including glucose-6-phosphate (M+6), 3-phosphoglycerate (3PG) / 2-phosphoglycerate (2PG) (M+3), phosphoenolpyruvate (PEP) (M+3) and pyruvate (M+3) declared significantly reduced glycolysis pathway by NMNH treatment. The results demonstrated that NMNH repressed cellular glycolysis. Meanwhile, the production of TCA cycle intermediates from glycolysis was clearly suppressed by NMNH based on the results of citrate (M+2), α-ketoglutarate (M+2), succinate (M+2) and malate (M+2) in Figure S3D. However, NMN treatment revealed distinct regulation of glycolysis and TCA cycle. NMN only induced mild decrease of 2PG/3PG and PEP in glycolysis (Figure S3C) and almost no observable reduction of TCA cycle in cells (Figure S3B). These results suggested that NMNH has very significant suppression of glycolysis and TCA cycle while NMN also hindered the glycolysis but with much milder extent.

### NMNH Repressed Cell Growth

Since glycometabolism is very important to cell growth while NMNH repressed glycolysis and TCA cycle in HepG2 cells, we explored the effect of NMNH on cell growth and found that NMNH treatment inhibited HepG2 cell growth at concentrations higher than 250 μM (Figure 4A). We performed data dependent quantitative proteomic analysis to identify differentially expressed proteins (DEPs) of HepG2 under 1 mM NMNH treatment for 12 h. 6142 proteins were identified using biological triplicates with false-positive rate less than 1%. The fold change cutoff was determined according to 88% coverage of percentage variations (Figure S4A) and proteins with the fold change more than 1.3 or less than 0.77 were considered as DEPs.^17, 18^ Based on tandem mass tag (TMT) ratio in proteins with 2 or more unique peptides, we identified 289 up-regulated proteins and 171 down-regulated proteins (Figure S4B). Ingenuity Pathway Analysis (IPA) revealed that NMNH treatment affected cell cycle progression (Figure S4C). We analyzed the cell cycle in HepG2 cells treated with 1 mM NMNH or NMN for 12 h, respectively and uncovered that NMNH caused cell cycle arrest whereas NMN had no effects on cell cycle (Figure 4B). The representative histograms and gating strategies are shown in Figure S4D. To further examine the effects of NMNH on cell growth, we treated 786-O cells with different concentrations of NMNH. 786-O cells, derived from a clear cell renal cell carcinoma (ccRCC) patient, exhibit a typical Warburg phenotype and their growth is dependent on glycolysis.^19, 20^ We measured the growth curve of 786-O cells and found that 500 μM NMNH completely blocked cell growth (Figure 4C). Further analysis revealed that NMNH inhibited cell growth in 786-O cells at 50 μM (Figure 4D), whereas 250 μM NMNH was needed to effectively inhibit cell growth in HK-2 cells (Figure 4D), which was an immortalized proximal tubule epithelial cell line from normal adult human kidney.^21^

**Figure 4.**
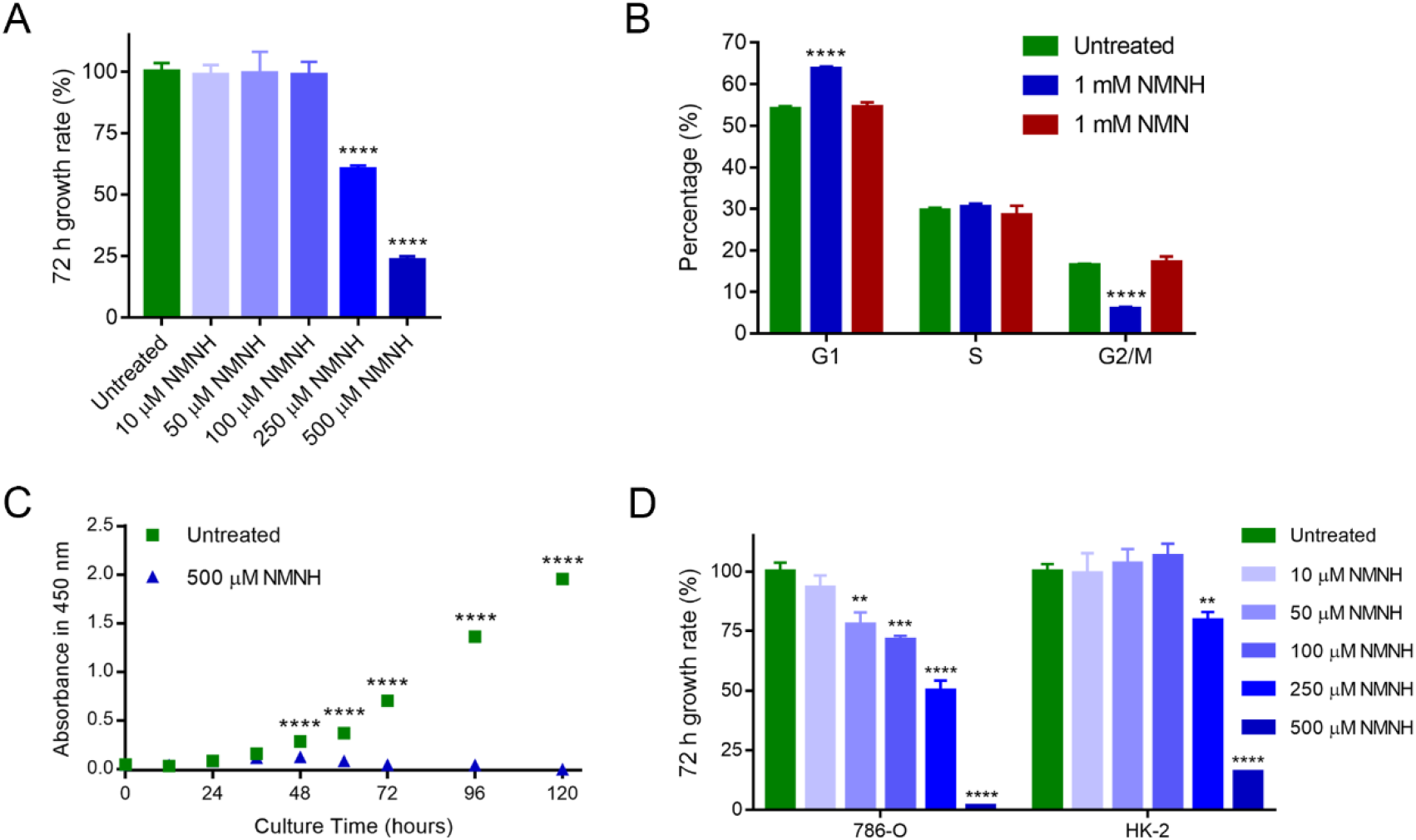
NMNH repressed cell growth *in vitro*. (A) High concentration NMNH inhibited the growth of HepG2. Cell growth rates were determined after NMNH treatment for 72 h using CCK-8 (Dojindo, Kumamoto, Japan). Data are shown as mean ± SD (n = 3). ****p < 0.0001. Significance is obtained compared to the untreated group unless otherwise specified. (B) Percentages of G1phase, S phase, and G2/M phase subpopulations in differently treated HepG2 cells by cell cycle analysis. Data are shown as mean ± SD (n = 5). ****p < 0.0001. Significance is obtained compared to the untreated group unless otherwise specified. (C) Growth curve of 786-O under 500 μM NMNH treatment. Cell growth rate were determined under 500 μM NMNH treatment using CCK-8 (Dojindo, Kumamoto, Japan). Data are shown as mean ± SD (n = 3). ****p < 0.0001. Significance is obtained compared to the untreated group unless otherwise specified. (D) 786-O is more sensitive to NMNH compared to HK-2. Cell growth rates were determined after NMNH treatment for 72 h using CCK-8 (Dojindo, Kumamoto, Japan). Data are shown as mean ± SD (n = 3). **p < 0.01, ***p < 0.001, ****p < 0.0001. Significance is obtained compared to the untreated group unless otherwise specified.

### NMNAT catalyzed NAD^+^ biosynthesis from NMNH

Our results demonstrated that the cellular NAD^+^ and NADH levels were elevated prominently by NMNH. In order to investigate the specific mechanism of NMNH to enhance the biosynthesis of NAD^+^ and NADH, the isotope tracing was performed using ^13^C_6_-glucose. The tracer of ^13^C_6_-glucose enables to provide the ribosyl labeling either through NMN from PRPP or ATP from purine biosynthesis (Figure S5A). Four different labeling status were monitored using TSQ mass spectrometer as shown in Figure 5A. Interestingly, unlike TCA cycle regulation, NMN and NMNH treatments illustrated similar trend of NAD^+^ and NADH biosynthesis reprogramming (Figure 5B and 5C). The incorporation of exogenous NMN or NMNH stimulated the biosynthesis of NAD^+^ and NADH, which resulted in the enhanced labeling from ATP (M+5-A). However, NMNH had more striking effect to stimulate NAD^+^ and NADH biosynthesis with almost completely inhibiting endogenous NMN synthesis by nicotinamide phosphoribosyltransferase (NAMPT), as minimal percentage of ribosyl labeled (M+5-N and M+10) NAD^+^ and NADH were detected.

**Figure 5.**
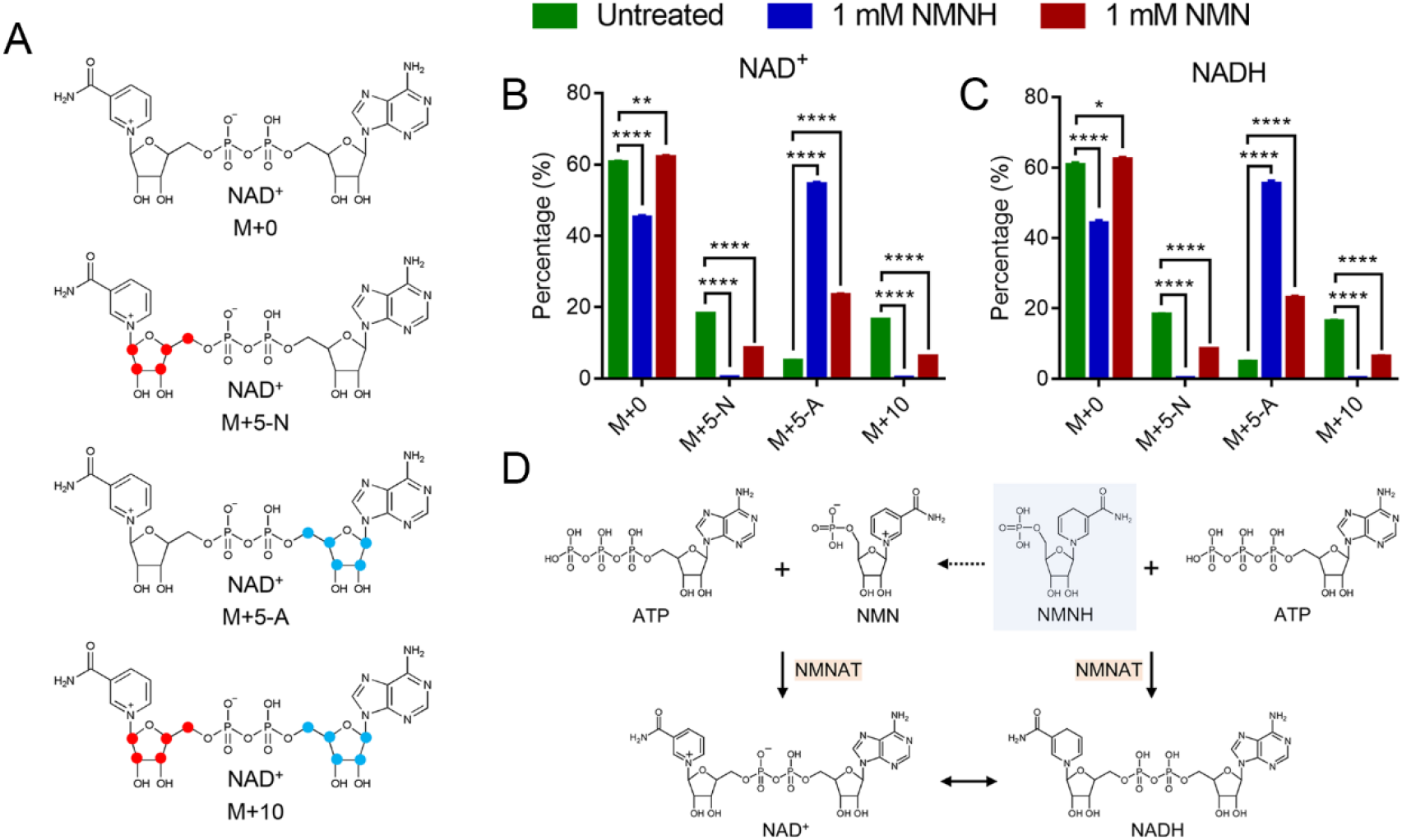
NMNH-induced NAD^+^ synthesis pathway deconstructing. (A) NAD^+^ label patterns. Red dots and blue dots represent ^13^C atoms. (B) Percentages of different isotope-encoded NAD^+^. M+*n* represents NAD^+^ contains *n* ^13^C atoms. Cellular NAD^+^ concentrations were determined after 1 mM NMN or NMNH treatment in media supplied with U-^13^C_6_ glucose for 6 h using TSQ Quantiva mass spectrometer. Data are shown as mean ± SD (n = 4). **p < 0.01, ****p < 0.0001. Significance is obtained compared to the untreated group unless otherwise specified. (C) Percentages of different isotope-encoded NADH. M+*n* represents NADH contains *n* ^13^C atoms. Cellular NADH concentrations were determined after 1 mM NMN or NMNH treatment in media supplied with U-^13^C_6_ glucose for 6 h using TSQ Quantiva mass spectrometer. Data are shown as mean ± SD (n = 4). *p < 0.05, ****p < 0.0001. Significance is obtained compared to the untreated group unless otherwise specified. (D) Proposed metabolic pathway through which NMNH is synthesized to NAD^+^

The recent study suggested that NRH was phosphorylated to NMNH which was further converted to NADH by NMNAT for boosting cellular NAD^+^ content.^12^ Tannic acid^12^, which is a specific NMNAT inhibitor, was used to explore the contribution of NMNAT in NAD^+^ biosynthesis (Figure S5B). Meanwhile, the effect of FK866 was also examined as a potent NAMPT inhibitor. The results suggested that tannic acid almost abolished the NAD^+^ increasing effect of NMNH while FK866 only showed slightly inhibitory effect (Figure S5B). To further confirm that NMNAT catalyzed the NMNH to NAD(H), we constructed NMNAT1-knockdown cell lines in HepG2 cells by shRNA (Figure S5C) and found that NMNAT1 knockdown compromised the NAD^+^ enhancing effect of NMNH (Figure S5D).

It was also found that the cellular NMN level was increased by NMNH even more dramatically than NMN treatment. (Figure S5E). Besides, the isotope tracing results showed < 0.4% of ribosyl labeled NMN in NMNH treated cells (Figure S5F), which suggested that the increased NMN was not biosynthesized from NAM and PRPP. This gave the hint of NMN production from NMNH in cells. Former reports indicated that ribosyldihydronicotinamide dehydrogenase (NQO2) could use NRH or a variety of its analogues as electron donors to reduce quinones to generate NR or relative dehydrogenation products.^22, 23^ It provided the possibility that NQO2 may have the function of turning NMNH to NMN. Herein, we constructed NQO2-knockdown cell lines in HepG2 cells by shRNA (Figure S5G). However, NQO2 knockdown didn’t alter the NMN level in NMNH-treated cells (Figure S5H), suggesting that NQO2 was not responsible for the conversion of NMNH to NMN although it contributed to the synthesis of NAD^+^. Further studies are needed to explore how NMNH was converted into NMN. Based on the results, we suggested that NMNH increased cellular NAD^+^ content in two ways that NMNH was directly converted to NADH or it was firstly converted to NMN and then to NAD^+^, as shown in Figure 5D.

## DISCUSSION

NAD^+^ is an important regulator of health and longevity.^24, 25^ Dozens of human clinical trials of NAD^+^ precursors are ongoing or recruiting participants currently while the existing data suggest that the translation from mouse model to human is not straightforward as people thought.^26^ Therefore, finding new agents that boost NAD^+^ levels is important to increase candidates for improving human health, especially for elderly people. Herein, we established a reduction method for efficient preparation of NMNH from NMN. Our study supports previous findings that the reduced form of NAD^+^ precursors (NMNH and NRH) are better NAD^+^ enhancers than their corresponding ones in oxidized form both *in vitro* and *in vivo.*^11, 12^ Meanwhile, we confirmed that long-term administration of NMNH is safe to mouse.

Yang *et al.* reported that NRH substantially increased NAD^+^/NADH ratio in cultured cells and in liver suggesting that NRH only mildly increased NADH levels.^11^ However, NMNH prominently increased cellular NADH levels, both in cell line and in mouse liver. NAD^+^ and NADH are an essential cellular redox couple and are inter-converted.^27^ NADH is the electron carrier in respiratory chain for ATP generation.^28^ Using metabolomics and isotope tracing analysis, we identified that NMNH suppressed glycolysis as well as the TCA cycle. We also demonstrated that NMNH repressed cell growth in *in vivo* experiment. Unlike previous studies showing that NRH had little effect on cell growth ^11, 12^, we found that NMNH treatment caused cell cycle arrest and inhibited cell growth even at a concentration of 100 μM.

We also deconstructed NAD^+^ synthesis pathway induced by NMNH. We found that NMNH effectively increased cellular NAD^+^ content and this process was mainly dependent on NMNAT. Moreover, we suggest that NMNH treatment inhibited NAD^+^ synthesis through NAMPT for little NAD^+^ (M+5-N) and NAD^+^ (M+10) were detected, while NMN treatment showed less effect. We further suggested that NMNH were converted to NMN to synthesize NAD^+^. Based on the above results, we suggest that NMNAT catalyzes NMNH into NADH, which is then converted to NAD^+^. NMNH is the precursor of NADH and NMNH treatment increases cellular NADH level. NAD^+^ has been well studied for its health benefits, while the biological effects of NADH are less known. A recent study indicated that NADH level was critical to maintain mice's fertility, which suggested that NADH may have distinct biological benefits that have been discovered.^29^ We suggest that NMNH is able to help people learn more about the biological functions of NADH.

In summary, we developed a chemical reduction method to produce NMNH in high yield. Our results demonstrate that NMNH is a potent NAD^+^ enhancer *in vitro* and *in vivo*. NMNH also significantly increased cellular NADH levels, repressed glycometabolism and inhibited cell growth.

## SIGNIFICANCE

NAD^+^ is an essential metabolite in living organisms and has been reported to be a health-promotion molecule. Here, we introduce a method to generate the reduced-form NAD^+^ precursor, NMNH, which is a more potent NAD^+^ enhancer than NMN both *in vitro* and *in vivo*. Besides, we found that NMNH can suppress glycolysis and TCA cycle, as well as repress cell growth. Contrast to NMN, which is the direct precursor of NAD^+^, NMNH is the direct precursor of NADH. Using proteomics and metabolomics technologies, our work demonstrated the effects of NMNH, which is a reduced-form NAD^+^ precursor, on cell metabolism. The significance of our findings is that we demonstrated a new potent NAD^+^ enhancer and explored the biological effects of NAD^+^ precursor in reduced form.

## AUTHOR CONTRIBUTIONS

Conceptualization, Y.L. and H.D.; Methodology, Y.L., C.L., T.L., W.Z., Z.Z., X.L. and H.D.; Investigation, Y.L., C.L. and T.L.; Validation, Y.L., C.L. and T.L.; Writing – Original Draft, Y.L. and T.L.; Writing – Review & Editing, Y.L., X.L. and H.D.; Funding Acquisition, H.D.; Supervision, H.D.

## ACKNOWLEGEMENT

We thank Beijing Advanced Innovation Center for Structural Biology and the Facility for Protein Chemistry and Proteomics,Metabolomics Facility, and the Cell Flow Cytometry Facility at Tsinghua University. We thank Michelle Goody, PhD, from Liwen Bianji, Edanz Editing China (www.liwenbianji.cn/ac), for editing the English text of a draft of this manuscript. This study was supported by the Chinese Ministry of Science and Technology (grant No. 2017ZX10201101 and No. 2020YFC2002705), China Postdoctoral Science Foundation (grant No. 2017M610080), National Key Research and Development Program (grant No. 2017YFA0505103), and National Nature Science Foundation of China (grant No. 21877068).

## METHODS

### NMNH preparation

NMNH was prepared by reducing NMN with TDO. Briefly, 340 mg NMN and 125 mg TDO were mixed in 1 mL 10% ammonia solution and incubated in 40°C water bath for 1 h. Next, NMNH was purified by a Dionex UltiMate 3000 (Thermo Scientific) HPLC system with an amide column (Waters, XBridge, 10 × 250 mm). Mobile phase A was prepared by adjusting HPLC-grade water to pH 10.0 using ammonia solution and mobile phase B was prepared by adjusting HPLC-grade acetonitrile to pH 10.0 using ammonia solution. Follow rate was set at 3 mL / min and the gradient was as follows: 0 min, 90% B; 5min, 90% B; 45 min, 40% B; 50 min, 40% B; 55 min, 90% B; 60 min, 90% B. The fraction with 340 nm absorption was collected and concentrated by vacuumdrying. NMNH was also generated by enzymatic decomposing NADH using NudC. Briefly, 20 mg NADH and 13 μg NudC were dissolved in 1 mL H_2_O containing 500 mM ammonium acetate and 3 mM magnesium chloride. The mixture was incubated in 48°C water bath for 1 h. Characteristic absorption feature and stability of NMNH were characterized using the same HPLC system, mobile phases and follow rate. The gradient was as follows: 0 min, 90% B; 5 min, 90% B; 15 min, 40% B; 18 min, 40% B; 19 min, 90% B; 20 min; 90% B.

### Cell lines

Human hepatocellular carcinoma cell line HepG2 (male), human renal cell adenocarcinoma cell line 786-O (male), human ovary clear cell carcinoma cell line ES-2, mouse embryo fibroblast cell line 3T3-L1 and human kidney cell line HK-2 (male) were purchased from the cell bank of Chinese Academy of Sciences (Shanghai, China). HepG2 and 786-O cells were cultured in RPMI1640 medium (Wisent, Canada) with 10% fetal bovine serum (PAN-Biotech, Germany) and 1% penicillin and streptomycin (Wisent, Canada) supplementation. ES-2 cells were culture in McCoy's 5A medium (Gibco) with 10% fetal bovine serum (PAN-Biotech, Germany) and 1% penicillin and streptomycin (Wisent, Canada) supplementation. 3T3-L1 cells were cultured in DMEM (Wisent, Canada) medium with 10% fetal bovine serum (PAN-Biotech, Germany) and 1% penicillin and streptomycin (Wisent, Canada) supplementation. HK-2 cells were cultured in Defined Keratinocyte SFM (Gibco) supplemented with 2.5 μg / 500 mL EGF recombinant human protein (Gibco) and 1% penicillin and streptomycin (Wisent, Canada). Cells were cultured in cell incubator containing 5% CO_2_ at 37 °C.

### *In vivo* experiments

Wild type C57BL/6J male mice, weighing 25±3g, 8 weeks old were used for experiment, which were randomly divided into three groups including PBS-treated group, NMNH treated group and NMN treated group. All the mice were housed in a temperature and light regulated room in a SPF facility and received food or water *ad libitum*. All animal experiments conform to the guidelines of the Laboratory Animal Research Center of Tsinghua University. All animal protocols used in this study were approved by the Institutional Animal Care and Use Committee of Tsinghua University. The sera were obtained by retro-orbital blood collection. The serum ALT and AST levels were measured in animal hospital of China Agricultural University using an automatic biochemical analyzer (Cobas C501, Roche).

### NudC expression and purification

DNA corresponding to the NudC open reading frame was synthesized (QINGLAN BIOTECH) and integrated into pET21b vector. Expression plasmid was transformed into C41 (DE3) competent cell (TianWeiTaiDa, China). Briefly, *E. coli* cells transformed with expressing plasmid were cultured in LB media until OD_600_ reached 0.6 and protein expression was induced by 1mM IPTG. Harvested cells were lysed by ultrasonication in lysis buffer (300 mM NaCl, 50 mM NaH2PO4, 10 mM imidazole, pH 8.0). Cell lysates were centrifuged at 15, 000×*g* for 30 min and the supernatants were purified by Ni-NTA agarose (Qiagen) and desalted using a HiTrap Desalting column (GE Healthcare).

### Mass spectrometric analysis of NMNH

NMNH products were analyzed by a Q Exactive mass spectrometer (Thermo Scientific™) connected to a Dionex UltiMate 3000 (Thermo Scientific) HPLC system without column in negative ion mode. The mobile phase A was prepared by adjusting HPLC-grade water to pH 10.0 using ammonia solution. Follow rate was set at 0.2 mL / min and gradient was as follows: 0 min, 100% A; 5min, 100% A. Detailed mass spectrometer parameters were set as follows: spray voltage was set at 3.5 kV; capillary temperature was set at 250°C; sheath gas flow rate (arb) was set at 45; aux gas flow rate (arb) was set at 10; mass range (m/z) was set at 80-750; full MS resolution was set at 70, 000; MS/MS resolution was set at 17, 500; topN was set at 10; HCD energy was set at 20, 30, 40.

### Establishment of stable NMNAT1 and NQO2 knockdown cell lines

The shRNA-containing plasmids (pLKO.1) for NMNAT1 and NQO2 knockdown were purchased form the shared Instrument facility at the center for biomedical analysis of Tsinghua University. The shRNA-containing plasmids were co-transfected with pLP2, pLP/VSVG and pLP1 into 293T cells by polyethylenimine. After 48 h, cell culture supernatants were collected and concentrated using PEG6000. The lentiviral particles were resuspended using PBS. HepG2 cells were transfected with lentiviral particles for 10 h in the presence of 10 μg/mL of polybrebe. Cells were selected under 2 μg/mL of puromycin to obtain stable knockdown cell lines which were then verified by western blot.

### Metabolomic analysis

Polar metabolites were extracted from cells and tissues (mouse liver) using pre-chilled 80% methanol (vol / vol) according to the method described before.^30, 31^ For cell samples, cells were washed for three times using cold PBS to avoid serum contamination and stored at −80 °C for 2 h after addition of 2 mL of 80% methanol. Cells were transferred into 1.5 mL tubes and the supernatants were harvested after centrifugation. The supernatants were dried by vacuum drying and stored −80 °C before analyzing. The pellets were dissolved using 1 M KOH for protein concentration determination. Metabolite samples were redissolved using 80% methanol according to protein concentration for mass spectroscopy analysis. For tissue samples, 25 mg tissues were homogenized in 250 μL pre-chilled 80% methanol and stored at −80 °C for 2 h. Samples were centrifuged and supernatants of the same volume were transferred into new 1.5 mL tubes. The supernatants were dried by vacuum drying and redissolved using the same volume of 80% methanol for mass spectroscopy analysis. Polar metabolites were analyzed by a Q Exactive Orbitrap mass spectrometer (Thermo, CA) or a TSQ Quantiva Ultra triple-quadrupole mass spectrometer (Thermo Fisher, CA). For Q Exactive Orbitrap mass spectrometer analysis, an Ultimate 3000 UHPLC (Dionex) was coupled to the mass spectrometer which was in positive mode. Atlantis HILIC Silica column (2.1×100 mm, Waters) was used for sample separation. In positive mode, mobile phase A was prepared by dissolving 0.63 g of ammonium formate in 50 ml of HPLC-grade water followed by adding 950 ml of HPLC-grade acetonitrile and 1 μl of formic acid. Mobile phase B was prepared by dissolving 0.63 g of ammonium formate in 500 ml of HPLC-grade water followed by adding 500 ml of HPLC-grade acetonitrile and 1 μl formic acid. The elution gradient was as follows: 0 min, 1% B; 2 min, 1% B; 3.5 min, 20% B; 17 min, 80% B; 17.5 min, 99% B; 19 min, 99% B; 19.1 min, 1% B; 22 min, 1% B. In negative mode, mobile phase A was prepared by 0.77 g of ammonium acetate in 50 ml of HPLC-grade water followed by adding 950 ml of HPLC-grade acetonitrile and pH was adjusted to 9.0 using ammonium hydroxide. Mobile phase B was prepared by dissolving 0.77 g of ammonium acetate in 500 ml of

HPLC-grade water followed by adding 500 ml of HPLC-grade acetonitrile and pH was adjusted to 9.0 using ammonium hydroxide. The elution gradient was as the same as positive mode. Detailed parameters of mass spectrometer were as follows: spray voltage was set at 3.5 kV; capillary temperature was set at 275°C; sheath gas flow rate (arb) was set at 35; aux gas flow rate (arb) was set at 8; mass range (m/z) was set at 70–1050; full MS resolution was set at 70,000; MS/MS resolution was set at 17,500; topN was set at 10; NCE was set at 15/30/45; duty cycle was set at 1.2 s.

For TSQ Quantiva Ultra triple-quadrupole mass spectrometer analysis, a Dionex Ultimate 3000 UPLC system was coupled to the mass spectrometer, equipped with a heated electrospray ionization (HESI) probe. Extracts were separated by a synergi Hydro-RP column (2.0×100mm, 2.5 μm, phenomenex). A binary solvent system was used, in which mobile phase A consisted of 10 mM tributylamine adjusted with 15 mM acetic acid in water, and mobile phase B of methanol. This analysis used a 25-minute gradient from 5% to 90% mobile B. Positive-negative ion switching mode was performed for data acquisition. Cycle time was set as 1 s. The resolutions for Q1 and Q3 are both 0.7 FWHM. The source voltage was 3.5 kV for positive and 2.5 kV for negative ion mode. The source parameters are as follows: spray voltage was set at 3.0 kV; capillary temperature was set at 320°C; heater temperature was set at 300°C; sheath gas flow rate was set at 35; auxiliary gas flow rate was set at 10. Data analysis and quantitation were performed by the software Tracefinder 3.1.

For cell metabolomic experiments, 100 μM or 1 mM NMNH or NMN were used for treatment and the cells were treated for 1 h, 6 h or 12 h before metabolomic analysis. For tissue metabolomic experiments, 340 mg/kg NMNH or NMN were used for treatment and the mouse were treated for 6 h before metabolomic analysis. The experimental details are described in each figure legend.

### Western blot assay

Cells were harvested and lysed using RIPA lysis buffer (Beyotime, China) supplemented with 1% protease inhibitor cocktail (MERCK, Germany). After centrifugation at 14, 000×*g* for 15 min at 4°C, supernatants were collected and protein concentrations were determined using BCA protein assay kit (Solarbio, Beijing, China). Proteins of equal amounts were separated using 12% SDS-PAGE gel and transferred onto PVDF membrane. Western blot assay followed a standard procedure. Anti-acetyl-lysine antibody was purchased from MILLIPORE (16-272), anti-actin antibody was purchased from Cell Signaling Technology (4970S), anti-NMNAT1 antibody was purchased from Proteintech (11399-1-AP), anti-NQO2 antibody was purchased from Abcam (ab181049); anti-rabbit antibody was purchased form Cell Signaling Technology (7074S) and anti-mouse antibody was purchased from Cell Signaling Technology (7076S).

### Total NAD(H) concentration measurement by kit

Intracellular total NAD(H) concentration or NADH content was measured using NAD^+^/NADH detecting kit (Beyotime, China) following manufacture’s instruction. Briefly, cell media were removed and cells were washed with PBS. Cold NAD^+^/NADH extraction buffer was added to cells for cell lysis. Then the lysis was centrifuged (12, 000×*g*, 10 min, 4 °C).For cellular total NAD(H) concentration measurement, cell lysis supernatants and NADH standards were added into 96-well plate and mixed with ethanol dehydrogenase working solution and incubated at 37 °C for 10 min. Color reagent was then added to the mix and the absorbance at 450 nm was measured. For cellular NADH concentration measurement, cell lysis supernatants and NADH standards were heated at 60 °C for 1 h and were then added into 96-well plate and mixed with ethanol dehydrogenase working solution and incubated at 37 °C for 10 min. Color reagent was then added to the mix and the absorbance at 450 nm was measured. Total NAD(H) concentration or NADH concentration was calculated according to standard curve. The concentration measurements were detected in triplicate and data were analyzed using Student’s t-test.

### Cell proliferation assay

Cells were seeded in 96-well plates at proper density and cultured according to experimental requirement. Cell proliferation rates were determined with a Cell Counting Kit-8 (CCK-8) (Dojindo Laboratories, Kumamoto, Japan). Briefly, media were removed and cells were washed with PBS and incubated with media containing 10% CCK-8 (vol / vol) at 37 °C for 2 h. 450 nm absorbance was measured to represent relative cell number. Cell proliferation assays were performed in triplicate and data were analyzed using Student’s t-test.

### Proteomic analysis

200 μg proteins were extracted using 8 M urea followed by reduction and alkylation. Then, proteins were digested with trypsin (Promega, Fitchburg, WI) for 14 hours at 37 °C. Peptides were then desalted and labeled by tandem mass tag (TMTsixplex^TM^, Thermo) according to manufacturer’s protocol. Next, samples were mixed, desalted and separated by HPLC and analyzed by Orbitrap Fusion™ Lumos™ Tribrid™ mass spectrometer (Thermo Scientific™). The MS/MS spectra were searched against the Uniprot human database (release date of October, 06, 2019, 20352 sequences) using the SEQUEST searching engine of Proteome Discoverer 2.1 software. The experiments were performed with three biological repeats.

### Cell cycle determination

HepG2 cells were harvested after 1 mM NMNH or NMN treatment for 12 h and washed with PBS containing 1% FBS. PBS were removed and cells were resuspended using 70% pre-chilled ethanol. Cells were fixed over night in 4 °C. Ethanol were then removed and cells were resuspended using PBS supplemented with 40 μg/mL propidium iodide (PI, Leagene, China) and RNaseA (TIANGEN, China). Cell cycle analysis was performed on a BD Calibur Cytometer (Becton Dickinson, NJ)

### Isotope tracing metabolomic analysis

RPMI1640 medium that was glucose-free (Gibco, Thermo Fisher Scientific, USA) was supplemented with 11 mM ^13^C_6_-glucose (Cambridge Isotope Laboratories, USA) for cell culture. HepG2 cells were culture in ^13^C_6_-glucose RPMI1640 media supplemented with 1 mM NMNH or NMN for 6 h before metabolomic analysis. An unlabeled culture was prepared for unlabeled metabolites identification by adding equal concentration of unlabeled glucose instead of ^13^C_6_-glucose. Sample preparation and analyzing method was described in “Metabolomic analysis” section. Isotope tracing metabolomic analysis was performed using a TSQ Quantiva Ultra triple-quadrupole mass spectrometer.

### Statistical analysis

GraphPad Prism 7.00 was used for statistical analysis. Data were shown as mean ± SEM and Student’s t test was employed to determine significant differences (*p < 0.05, **p < 0.01, ***p < 0.001, ****p < 0.0001). p values of < 0.05 were considered to be significant. Significance is obtained compared to the untreated group unless otherwise specified.

---

**Figure S1.**
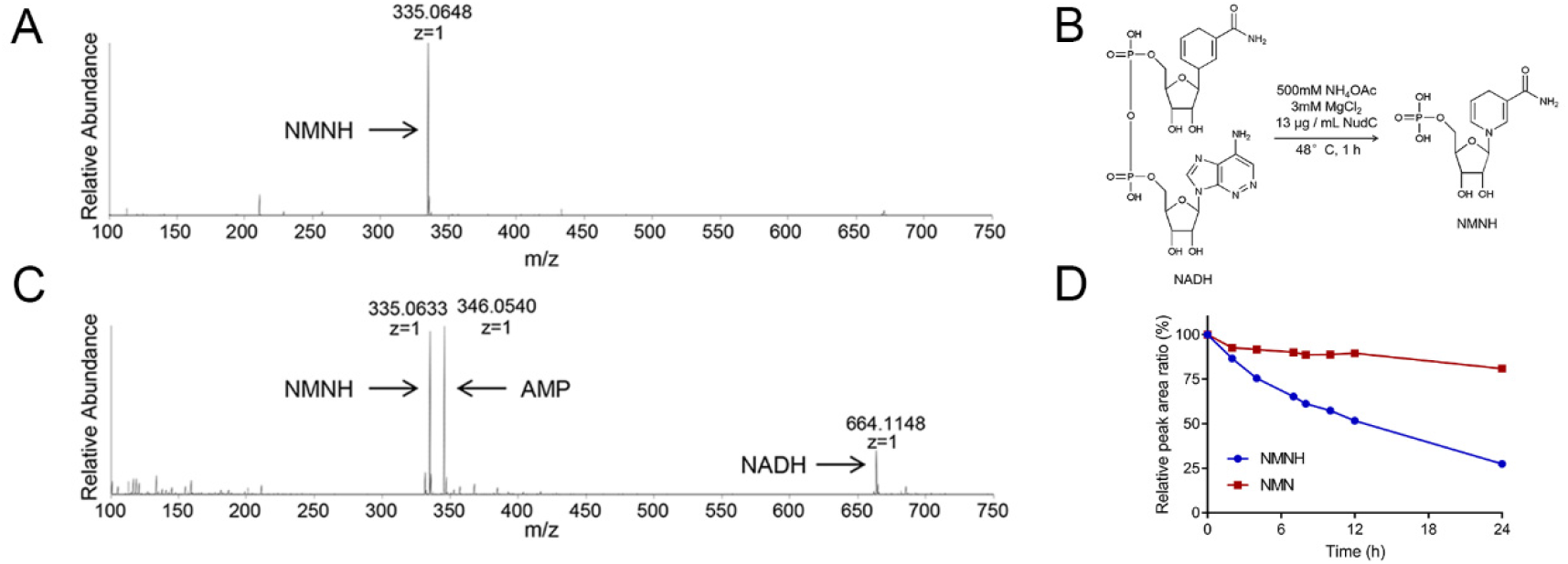
NMNH synthesis. (A) MS spectrum feature of NMNH generated by NMN reduction. NMNH has an m/z of 335.0648 in negative ion mode. (B) Procedure of NMNH generation by NADH decomposition. NMNH was generated by decomposing NADH using NudC. (C) MS spectrum feature of NMNH generated by NADH decomposition. NMNH has an m/z of 335.0633 in negative ion mode. (D) NMN is more stable than NMNH in cell medium. Equal amount of NMNH and NMN were dissolved in phenol red-free RPMI1640 medium with 10% fetal bovine serum and 1% penicillin and streptomycin supplementation and the 340 nm and 260 nm absorptions were determined.

**Figure S2.**
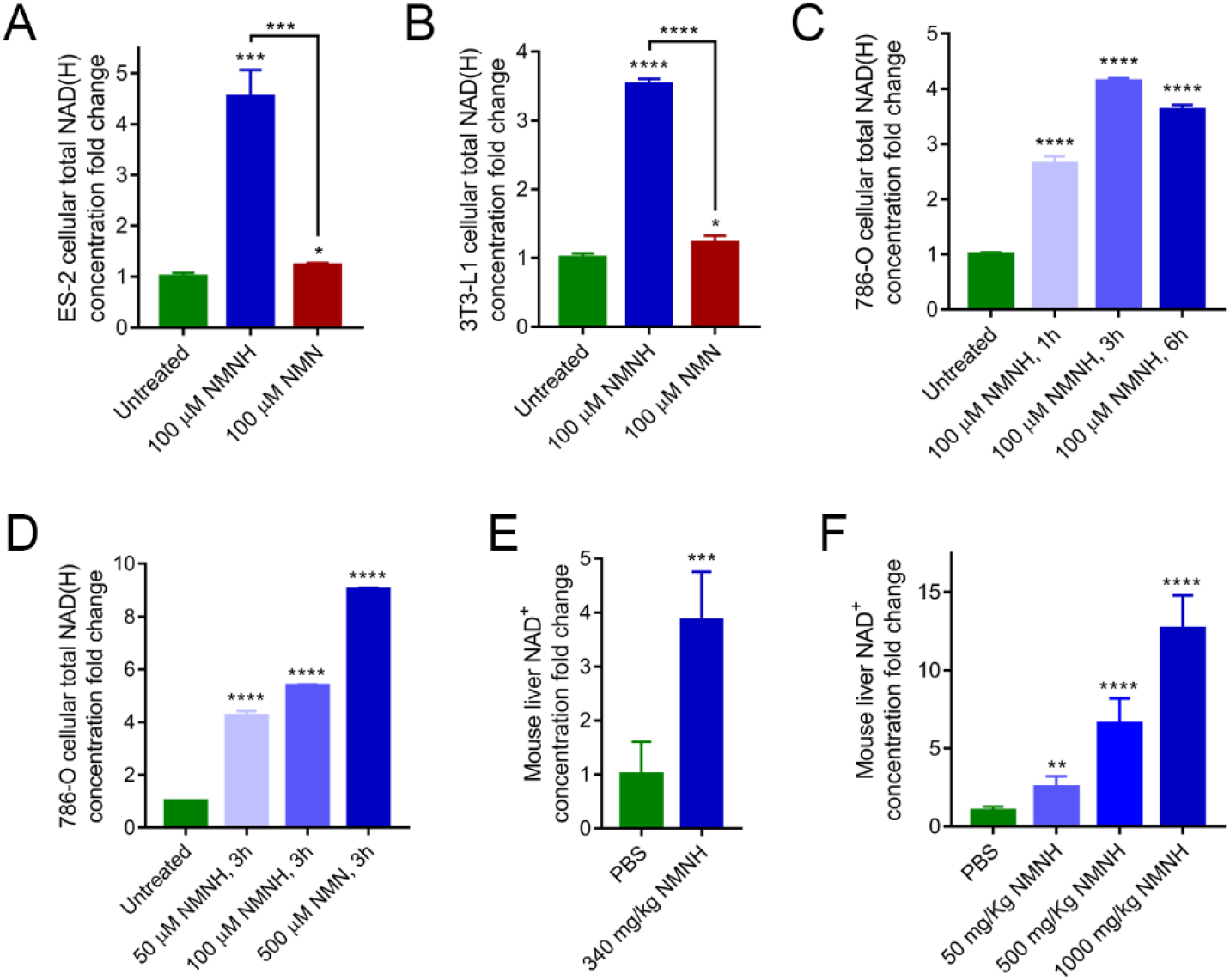
NMNH is potent NAD^+^ booster both *in vitro* and *in vivo*. (A) NMNH increased cellular total NAD(H) concentration in ES-2 cell. Cellular total NAD(H) concentration was determined after 100 μM NMN or NMNH treatment for 3 h using NAD^+^/NADH detecting kit (Beyotime, China). Data are shown as mean ± SD (n = 3). *p < 0.05, ***p < 0.001. Significance is obtained compared to the untreated group unless otherwise specified. (B) NMNH increased cellular total NAD(H) concentration in 3T3-L1 cell. Cellular total NAD(H) concentration was determined after 100 μM NMN or NMNH treatment for 3 h using NAD^+^/NADH detecting kit (Beyotime, China). Data are shown as mean ± SD (n = 3). *p < 0.05, ****p < 0.0001. Significance is obtained compared to the untreated group unless otherwise specified. (C) NMNH increased cellular total NAD(H) concentration in 786-O cell. Cellular total NAD(H) concentration was determined after 100 μM NMNH treatment for 1-6 h using NAD^+^/NADH detecting kit (Beyotime, China). Data are shown as mean ± SD (n = 3). ****p < 0.0001. Significance is obtained compared to the untreated group unless otherwise specified. (D) NMNH increased cellular total NAD(H) concentration in 786-O cell. Cellular total NAD(H) concentration was determined after 50-500 μM NMNH treatment for 3 h using NAD^+^/NADH detecting kit (Beyotime, China). Data are shown as mean ± SD (n = 3). ****p < 0.0001. Significance is obtained compared to the untreated group unless otherwise specified. (E) NMNH is orally bioavailable. C57BL/6J male mice were treated with 340 mg/kg NMN or NMNH for 6 h by oral gavage. Liver NAD^+^ concentration was determined using TSQ Quantiva mass spectrometer. Data are shown as mean ± SD (n = 5). ***p < 0.001. Significance is obtained compared to the untreated group unless otherwise specified. (F) NMNH increased liver NAD^+^ concentration in a dosage-dependent manner. C57BL/6J male mice were treated with 50, 100, 500, 1000 mg/kg NMNH or PBS every other day for 1 week by intraperitoneal injection. Liver NAD^+^ concentration was determined using Q Exactive mass spectrometer. Data are shown as mean ± SD (n = 5 for PBS, 50 and 500 mg/kg groups; n = 4 for 1000 mg/kg groups).

**Figure S3.**
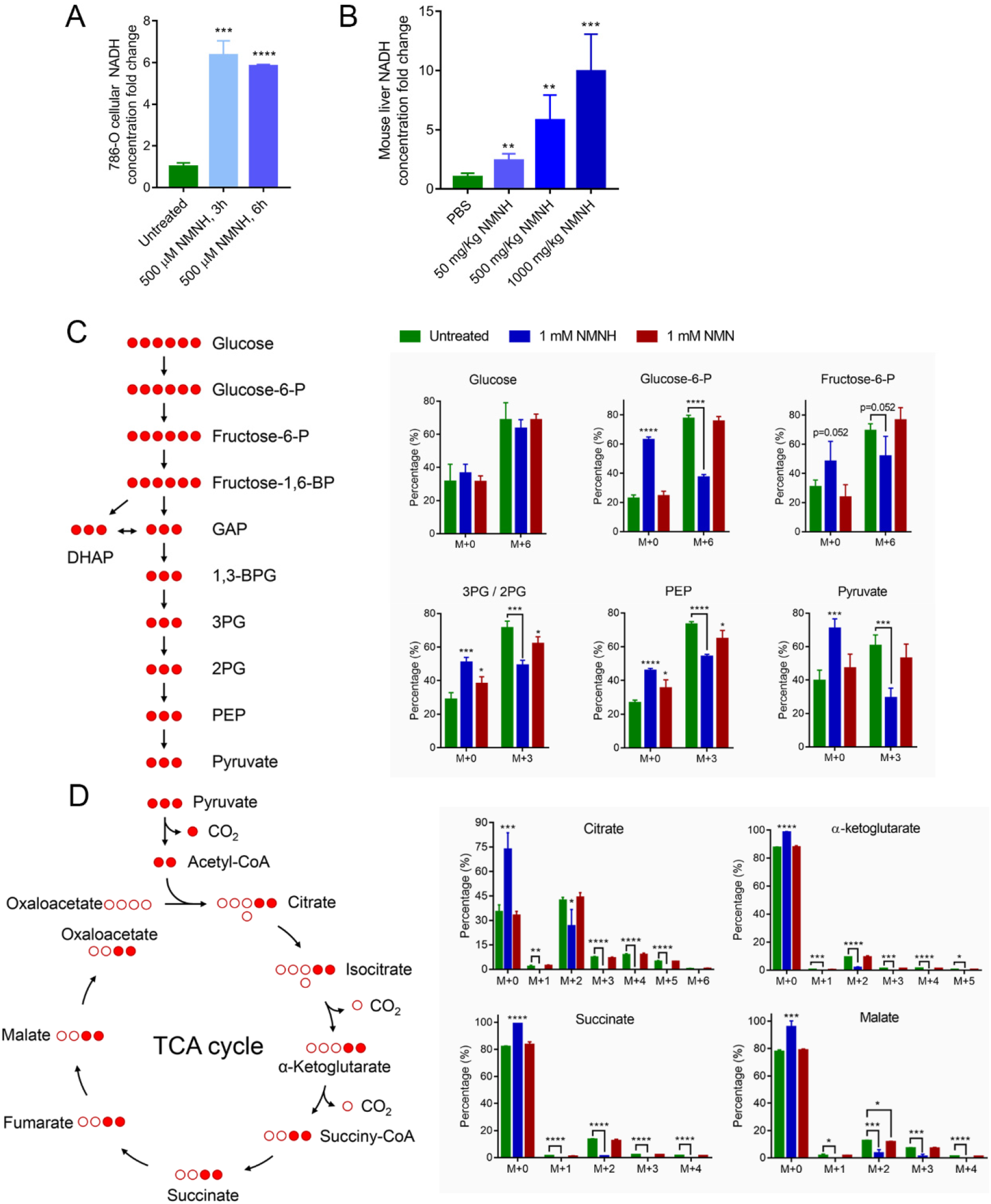
Metabolomic analysis of TCA intermediates under NMNH and NMN treatment. (A) 786-O cellular NADH concentration was determined after 500 μM NMN or NMNH treatment using NAD^+^/NADH detecting kit (Beyotime, China). Data are shown as mean ± SD (n = 3). ***p < 0.001, ****p < 0.0001. Significance is obtained compared to the untreated group unless otherwise specified. (B) NMNH increased liver NADH concentration in a dosage-dependent manner. C57BL/6J male mice were treated with 50, 100, 500, 1000 mg/kg NMNH or PBS every other day for 1 week by intraperitoneal injection. Liver NAD^+^ concentration was determined using Q Exactive mass spectrometer. Data are shown as mean ± SD (n = 5 for PBS, 50 and 500 mg/kg groups; n = 4 for 1000 mg/kg groups). (C) Left, schematic of glycolysis. Red solid circle represents ^13^C. Right, percentages of different isotope-encoded glycolysis metabolites. M+*n* represents a metabolite contains *n* ^13^C atoms. Cellular metabolite concentrations were determined after 1 mM NMN or NMNH treatment in media supplied with U-^13^C_6_ glucose for 6 h using TSQ Quantiva mass spectrometer. Data are shown as mean ± SD (n = 4). *p < 0.05, ***p < 0.001, ****p < 0.0001. Glucose-6-P, glucose-6-phosphate. Fructose-6-P, fructose-6-phosphate. Fructose-1,6-BP, fructose-1,6-bisphosphate. GAP, glyceraldehyde-3-phosphate. DHAP, dihydroxyacetone phosphate. 1,3-BPG, 1,3-bisphosphoglycerate. 3PG / 2PG, 3-phosphoglycerate / 2-phosphoglycerate. PEP, phosphoenolpyruvate. Significance is obtained compared to the untreated group unless otherwise specified. (D) Left, schematic of TCA cycle. Red solid circle represents ^13^C. Right, percentages of different isotope-encoded TCA cycle metabolites. M+*n* represents a metabolite contains *n* ^13^C atoms. Cellular metabolite concentrations were determined after 1 mM NMN or NMNH treatment in media supplied with U-^13^C_6_ glucose for 6 h using TSQ Quantiva mass spectrometer. Data are shown as mean ± SD (n = 4). *p < 0.05, **p < 0.01, ***p < 0.001, ****p < 0.0001. Significance is obtained compared to the untreated group unless otherwise specified.

**Figure S4.**
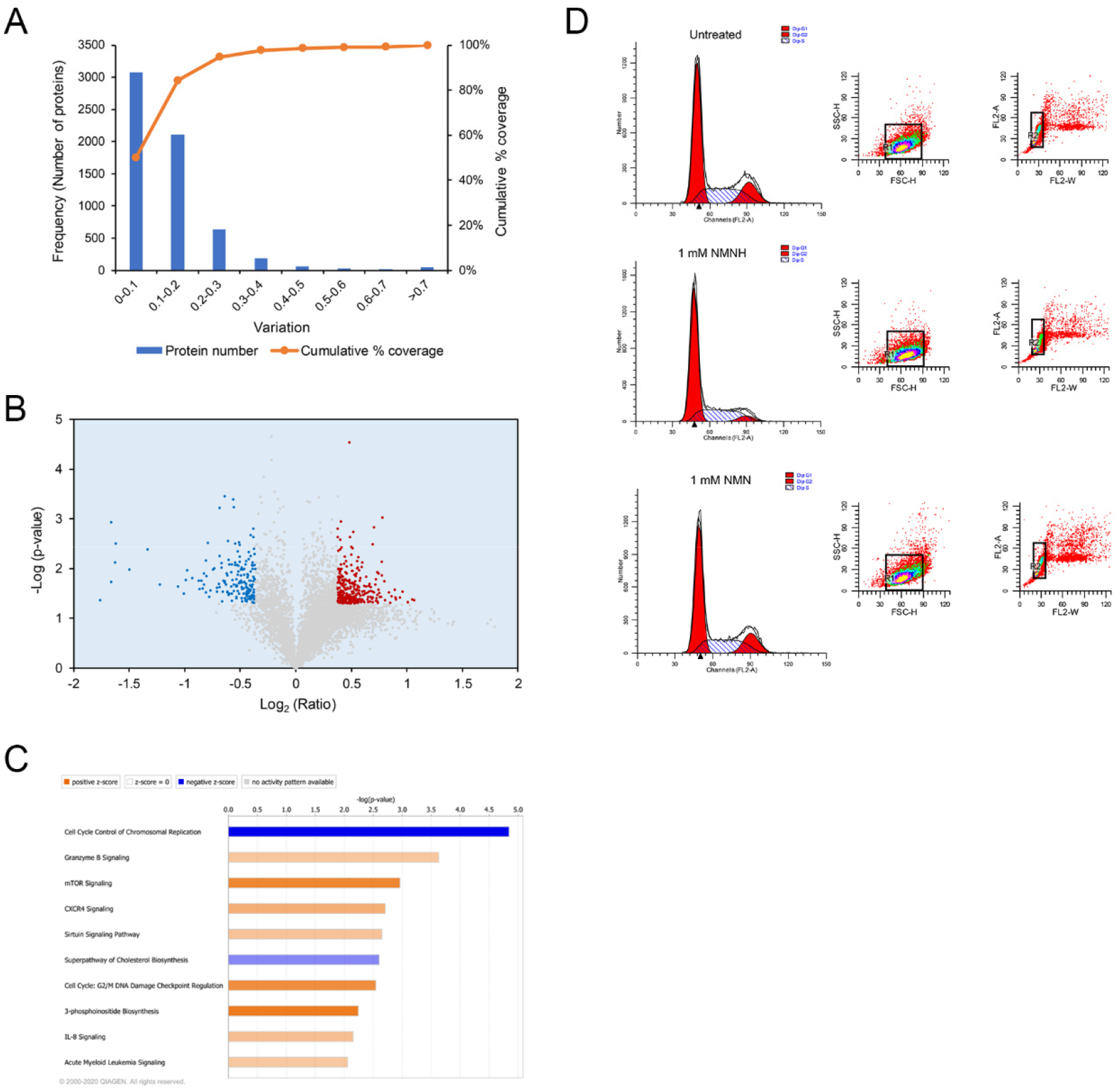
NMNH repressed cell growth *in vitro*. (A) Determination of experimental variation and cutoff value for the identified proteins. Horizontal axis represents TMT ratio variation. Primary vertical axis represents the number of proteins with corresponding different variation. Secondary vertical axis represents the cumulative percentage coverage of the counted proteins. Cutoff value was set according to the variation against 88% coverage of population. (B) Volcano plot obtained from TMT-based quantitative proteomics analysis. Red dots represent proteins exhibiting significative un-regulated fold changes. Blue dots represent proteins exhibiting significative down-regulated fold changes. (C) Representative canonical pathways enriched IPA (Ingenuity Pathway Analysis) software. (D) Representative histograms and gating strategies of cell cycle analysis. Upper panel, untreated group; middle panel, NMNH treated group; lower panel, NMN treated group.

**Figure S5.**
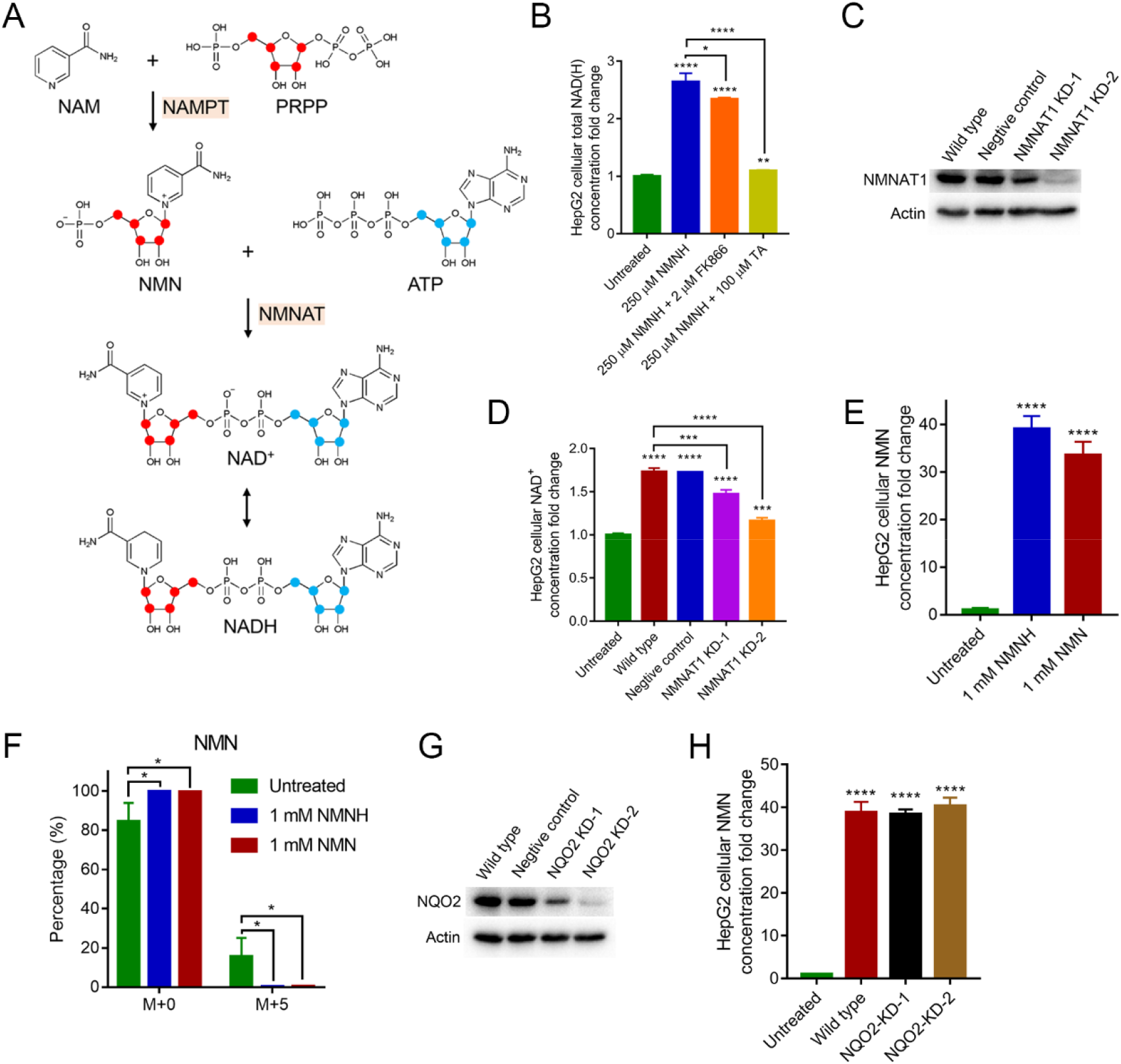
NMNH-induced NAD^+^ synthesis pathway deconstructing. (A) Schematic representation of the salvage pathway. Red dots and blue dots represent ^13^C atoms. (B) NMNH increased cellular NAD^+^ levels depending on NMNAT. Cells were treated by different kinds of inhibitors for 1 hour before NMNH treatment. Cellular total NAD(H) concentration was determined after 250 μM NMNH treatment for 1 h using NAD^+^/NADH detecting kit (Beyotime, China). Data are shown as mean ± SD (n = 3). *p < 0.05, **p < 0.01, ****p < 0.0001. Significance is obtained compared to the untreated group unless otherwise specified. (C) Western blot analysis of NMNAT1 expression in wild type, negative control and NMNAT1 knockdown HepG2 cells. (D) NMNAT1 knockdown compromised the synthesis of NAD^+^ from NMNH in HepG2 cells. Cellular NAD^+^ concentration was determined after 100 μM NMNH treatment for 1 h using Q Exactive mass spectrometer. Data are shown as mean ± SD (n = 4). **p < 0.01, ***p < 0.001, ****p < 0.0001. (E) NMNH increased cellular NMN concentration in HepG2 cell. Cellular NMN concentration was determined after 1 mM NMN or NMNH treatment for 6 h using Q Exactive mass spectrometer. Data are shown as mean ± SD (n = 4). ****p < 0.0001. (F) Percentages of different isotope-encoded NMN. M+*n* represents NMN contains *n* ^13^C atoms. Cellular NMN concentrations were determined after 1 mM NMN or NMNH treatment in media supplied with U-^13^C_6_ glucose for 6 h using Q Exactive mass spectrometer. Data are shown as mean ± SD (n = 4). *p < 0.05. (G) Western blot analysis of NQO2 expression in wild type, negative control and NQO2 knockdown HepG2 cells. (H) NQO2 knockdown didn’t alter the NMN level in NMNH-treated cells. Cellular NMN concentration was determined after 100 μM NMNH treatment for 1 h using TSQ Quantiva mass spectrometer. Data are shown as mean ± SD (n = 4). ****p < 0.0001.

